# Revisit the Inhibitory Effects of Glucocorticoids on Immunocytes

**DOI:** 10.1101/2023.01.28.525640

**Authors:** Shuting Wu, Shushu Zhao, Yiwei Zhong, Bin Wang

## Abstract

Glucocorticoids (GCs) are efficacious agents for reducing inflammation and suppressing immune responses, exerting various effects on immune cells through the intracellular glucocorticoid receptor (GR), and impacting both innate and adaptive immunity. In the context of COVID-19, glucocorticoids are often used to treat severe cases of patients by reducing inflammation, suppressing immune responses, and ameliorating the severity of COVID-19. However, the precise inhibitory effects on immune cells have yet to be comprehensively delineated. In this study, we extensively examined the inhibitory effects of treating Balb/c mice with dexamethasone (DEX) on lymphoid and myeloid cells. We observed that high doses of DEX treatment resulted in a reduction in the number of immunocytes and an attenuation of their activity. Particularly noteworthy, macrophages, DC cells, and monocytes were diminished by approximately 90% following high doses of DEX, while B cells experienced a reduction of about 70% and CD3 T cells were less affected. Furthermore, our findings demonstrated that DEX induces the inhibition of immune cells by engaging in high-affinity binding to GR. Consequently, we conclude that DEX treatments affect a broad range of immune cells, encompassing both lymphoid and myeloid cells, through depletion or the down-regulation of immune function, potentially acting via the GR signaling pathway. These findings may enhance the clinical applicability of DEX in achieving transient immune deficiency.

## Introduction

Glucocorticoids (GCs) are known inhibitors of the immune system. The synthesis and release of GCs are under dynamic circadian and ultradian regulation by the hypothalamic-pituitary–adrenal (HPA) axis[1]. Multiple studies have shown that GCs have potent anti-inflammatory and immunosuppressive properties[2–5]. Natural and synthetic glucocorticoids remain at the forefront of anti-inflammatory and immunosuppressive therapies used in the treatment of rheumatoid arthritis, multiple sclerosis, psoriasis, and eczema. Additionally, they are used in the treatment of certain leukemias and to combat the side effects of some chemotherapies and immunosuppressive regimens following organ transplant[4, 6, 7]. GCs induce large-scale lymphodepletion that differs from specific depletion mediated by antibodies in that it occurs predominantly through binding to the glucocorticoid receptor (GR) of immunocytes. GCs form a complex with the intracellular GR, which induces a conformational change in the receptor that allows translocation to the nucleus, where it binds to chromatin. This binding to chromatin leads to the regulation of gene expression, resulting in various anti-inflammatory and immunosuppressive effects [8–10]. In addition to genomic mechanisms, GCs elicit rapid effects by interacting directly with enzymes and other cell proteins, including the regulation of signaling pathways[11]. These rapid effects of GCs make them a promising treatment option for COVID-19, as they can help reduce the hyperactive immune response that often leads to severe lung inflammation. Dexamethasone (DEX), a widely used GC, has been shown to effectively achieve immune suppression and alleviate over-reactive inflammation in the lungs of COVID-19 patients. This treatment approach can be particularly beneficial for severe cases of the disease, where an excessive immune response plays a major role in worsening the patient’s condition[12]. However, the precise mechanism by which DEX interacts with immune cells and exerts its anti-inflammatory and immunomodulatory effects in this specific context is not fully understood. It is believed that DEX acts by inhibiting the production of pro-inflammatory molecules such as cytokines and chemokines, thereby reducing the recruitment and activation of immune cells in the lungs. Additionally, DEX may also interfere with the signaling pathways involved in immune cell activation, further suppressing the hyperactive immune response. Furthermore, the administration of DEX to DC results in a diminished level of major histocompatibility complex (MHC) class II and costimulatory molecules, consequently diminishing their capacity to activate T cells[13], while concurrently inhibiting the signal transmission pathway mediated by the T-cell receptor (TCR)[14]. Some reports have documented that GCs modulate apoptosis by repressing the expression of the Bcl-2 protein, which is encoded by the B-cell lymphoma/leukemia-2 gene[15]. Initial studies on the effects of GCs on B cells have yielded conflicting results, particularly with regards to the dosage relationship between GCs and B cell activity as well as antibody synthesis or levels[16]. GCs have also been found to suppress innate immune cells by regulating monocyte-macrophages. GCs can attenuate the exaggerated activitiy of immune cells that can arise during inflammation[17]. Intriguingly, glucocorticoids have been shown to enhance the accumulation and survival of neutrophils by inducing the expression of anti-apoptotic proteins such as members of the Bcl-2 family [18], which has contradictory effects on B cells. The regulatory effects of GCs or DEX on the immune system are intricate and still needed to fully understand their comprehensive impact on their effects on innate immune cells such as dendritic cells and monocyte macrophages, as well as their role in adaptive immune cells such as B and T cells. By examining all these aspects, a better understanding of these mechanisms will be crucial in optimizing the use of DEX as a treatment for severe cases of COVID-19 and other virally induced lung inflammations. In this study, we investigated the effects of DEX on the myeloid and lymphoid systems to reveal if the effects of DEX varied across different immune cell types. The effects on monocytes, DCs, macrophages, and NKTs were contingent on the dosage and duration, although CD3 T cells were less affected by the DEX treatments, whereas an enhanced level of neutrophils was evidence. These inhibitions by DEX exhibit a strong affinity for GR in the affected immune cells, suggesting a mechanistic basis for its immunosuppressive properties.

## Materials and Methods

### Mice and cells lines

Female Balb/c mice were housed and treated according to the guidelines of the Institutional Animal Care and Use Committee of the Fudan University in China. All mice were 6–8 weeks old and age-matched. All experiments were approved by the Committee of Experimental Animals of SHMC with the reference number 202012037S.

### Reagents

Dexamethasone (DEX) was purchased from Xi’an Reyphon Pharmaceutical CO (Xi’an, China). Mifepristone (RU486) was purchased from MCE (NJ, USA). Fluorochrome labeled antibodies against CD45(30-F11), CD4(GK1.5), CD8(53-6.7), CD25(3C7), CD11b(M1/70), F4/80, Ly6C(HK1.4), CD49b(DX5), B220(RA3-6B2) were purchased from BioLegend (CA, USA), CD11c(HL3), Ly6G(1A8) were purchased from BD Bioscience (CA, USA) and CD3 (145-2C11), Foxp3 (FJK-16S), fixable viability dye eFluor 780, Foxp3 transcription factor staining buffer kit were from eBioscience (OR, USA).

### Treatments

The mice were treated with 100 µg DEX in phosphate-buffered saline (PBS)/animal or vehicle by intraperitoneal injections for 3 consecutive days. Peripheral blood mononuclear cells (PBMC) were collected 24 h after the last administration of DEX. When determining the effect of dexamethasone dosage on immune cells, the doses per mouse were adjusted to 5 mg/kg for high-dose and 1.25 mg/kg for low-dose. Eight micrograms of RU486 were dissolved in 20 µl dimethyl sulfoxide (DMSO), added in 200 µl corn oil (MCE, NJ, USA), and vortexed. Mice were given 400 mg/kg body weight RU486 in corn oil intraperitoneally every 12 h before each DEX administration, and PBMC were collected 72 h after those treatments for FCM analysis (Supplementary 1A).

### Flow cytometry

For flow cytometry (FCM), whole blood with anticoagulant was treated with red cell lysis buffer. Single-cell suspensions were stained with viability dye eflour780 in PBS for 15 minutes at room temperature and then washed twice with PBS supplemented with 2% FBS. For detection of cell surface antigens, cells were stained with fluorochrome-tagged antibodies for 15 minutes at room temperature; for detection of Foxp3, cells were permeabilized and fixed using a commercial transcription factor staining buffer set (eBioscience, OR, USA). All stained samples were analyzed on an LSR Fortessa (BD, CA, USA) with FlowJo software (BD, CA, USA).

### Statistical analysis

The student’s t-test was used to compare continuous variables. Data were analyzed using one-way ANOVA for single-factor analysis among multiple groups and two-way ANOVA for multiple-dimensional comparisons. Data are shown as mean ± SEM or as percentages. P values reporting statistical significance were calculated in Prism 9 (GraphPad Software). P values less than 0.05 were deemed significant.

## Results

### Immunosuppressive Effects of Dexamethasone on Lymphoid and Myeloid Cells

In order to assess the impact of DEX on myeloid-derived innate immune cells, Balb/c mice were subjected to daily treatment with either 5 mg/kg DEX or vehicle (PBS) for a duration 3 days via the intraperitoneal (*ip*) route (Fig. 1A). Flow cytometry analysis of PBMC collected on the 4^th^ day (sFig. 1A and sFig. 2A) demonstrated a significant reduction in the levels of these cells, except for the neutrophils (Figs. 1B, 1C, and sFig. 1B). Further examination revealed that DEX induced the most substantial reductions in the proportions of macrophages (95%), DCs (90%), while demonstrating a comparatively smaller reduction in NK and NKT cells (Fig. 1D). Interestingly, the level of neutrophils was found to be increased, suggesting the involvement of a distinct regulatory mechanism. Given the relatively short half-life of DEX in circulation (2-3 days), we sought to explore whether its effects could persist after cessation of treatment. After 21 days of daily administration, the administrations were halted, and PBMC were analyzed. The data demonstrated that the proportions of the aforementioned affected myeloid-derived innate immune cells returned to normal levels 5 days after discontinuation of the drug (Figs. 2A-2F).

**Fig. 1.**
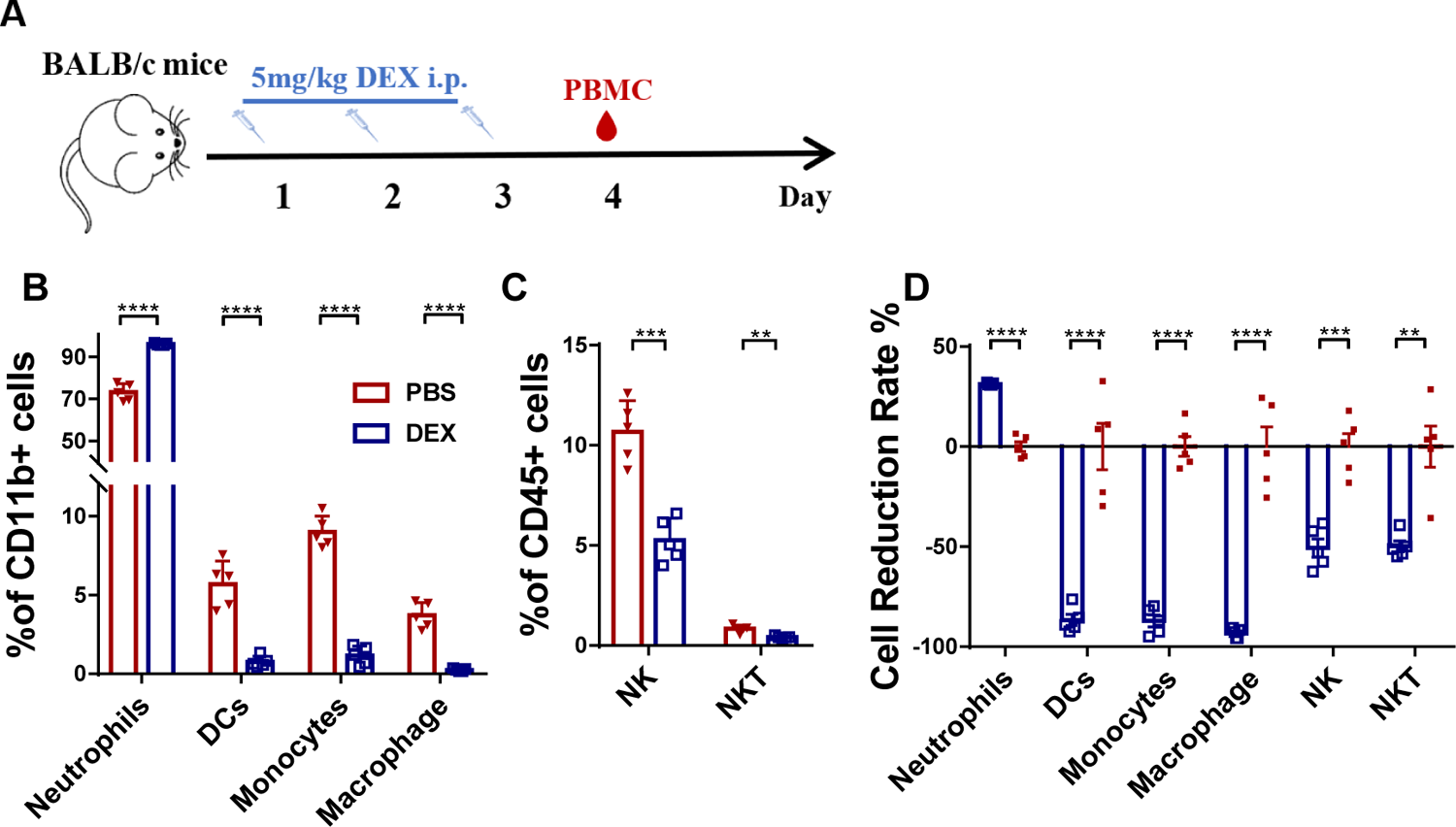
DEX’s immunoregulatory effects on myeloid-derived innate immune cells (A) Diagram of DEX treatment of mice. PBMC were isolated at 96 h after treatment with DEX and analyzed by flow cytometry (**B, C)**. Antibodies to Ly6G, CD11c, F4/80, Ly6C, and CD49b were used to identify neutrophils, DC, macrophages, monocytes, and NK cells. Antibodies to CD3 and CD49b were used to identify NKT cells. The results are presented in percentages of CD11b^+^ cells (**B**) and CD45^+^ cells (**C**); (**D**) shows the proportion of the decline in each innate immune cell type. Error bars indicate ± SEM, **** P < 0.0001, n = 5. Statistical significance was determined with an unpaired two-tailed T-test. Data are representative of two independent experiments.

**Fig. 2.**
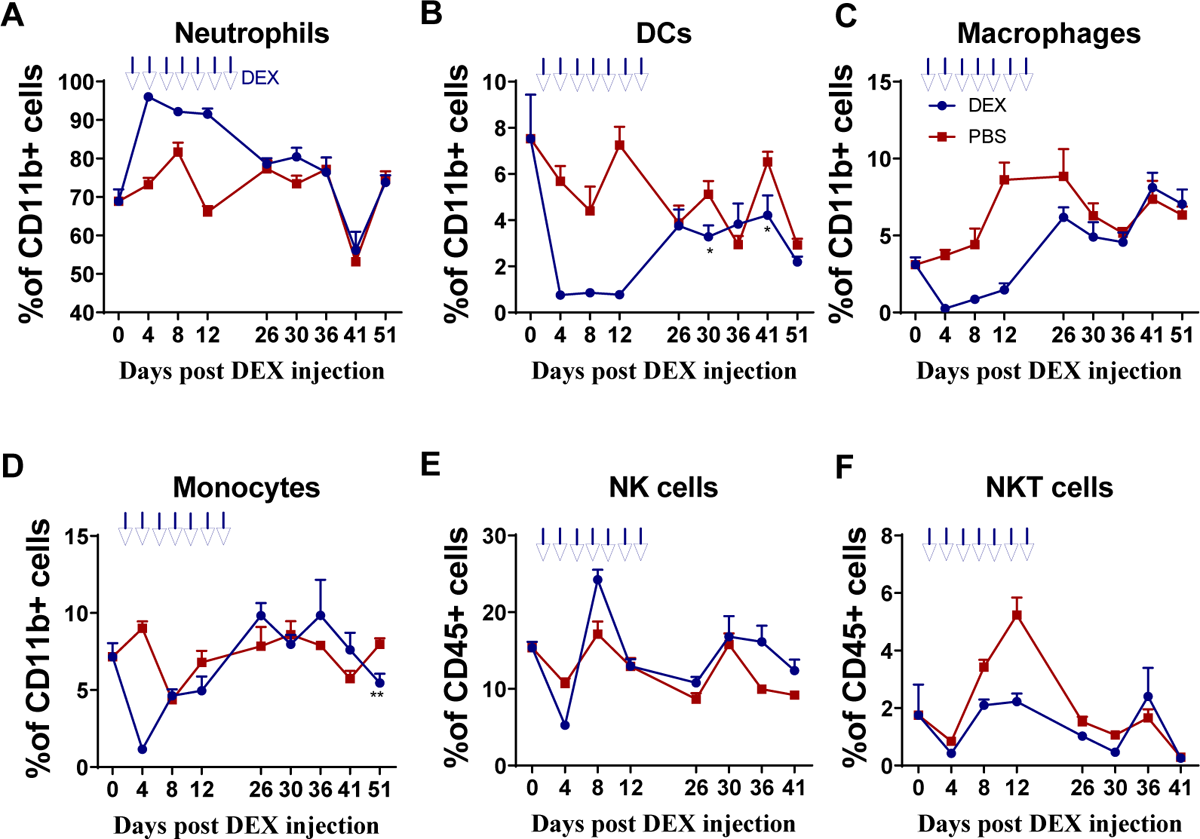
Recovery curves of innate immune cells after drug withdrawal PBMC were isolated on days 5, 9, 15, 20, and 30 after DEX withdrawal and analyzed by flow cytometry. Antibodies to Ly6G, CD11c, F4/80, Ly6C, and CD49b were used to identify neutrophils, DC cells, macrophages, monocytes, and NK cells; antibodies to CD3 and CD49b were used to identify NKT cells. Results from the DEX treatment group are shown in blue and from PBS vehicle group in red. Shown are the percentages of CD11b^+^ neutrophil cells (**A**), CD11b^+^ DC cells (**B**), CD11b^+^ macrophages (**C**), CD11b^+^ monocytes (**D**), CD45^+^ NK cells (**E**), and CD45^+^ NKT cells (**F**) Error bars indicate ± SEM, **** P < 0.0001. Statistical significance was determined with an unpaired two-tailed T-test. Data are representative of two independent experiments.

### Immunosuppressive Effects of Dexamethasone on Lymphoid-Derived Cells

The subsequent investigation aimed to analyze the immunosuppressive effects of DEX on lymphoid-derived cells, with a particular focus on T and B cells. The same treatment schedule as shown in Figure 1A was followed on Balb/c mice, and flow cytometry analysis was conducted to evaluate any alterations in CD3^+^, CD4^+^, CD8^+^, CD4^+^CD25^+^ T cells, and B220^+^ B cells (sFigs 1A and 2A). The results revealed that all analyzed cells were suppressed by DEX (Fig. 3A and sFig. 2B), with reductions of 72%, 40%, 45%, 40%, and 60%, respectively (Fig. 3B). Furthermore, the duration of the inhibitory effect on each of these cell types following drug withdrawal was assessed. Unlike myeloid-derived cells, lymphoid-derived cells exhibited a longer recovery period, taking up to 30 days to regain normal levels after cessation of treatment (Figs. 4A-4E).

**Fig. 3.**
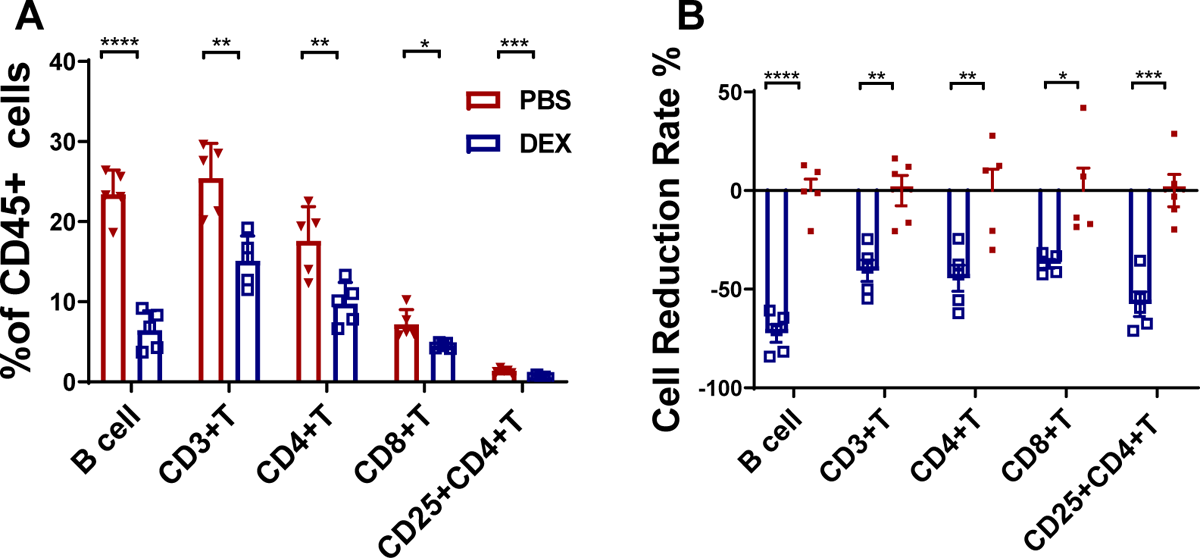
DEX’s immunoregulatory effects on lymphatic cells PBMC were isolated 96 h after treatment with 5 mg/kg DEX daily and analyzed by flow cytometry. Antibody to B220 was used to identify B cells. The results are presented as percentages of CD45^+^ cells (**A**) and the percentage of the decline in each lymphatic immune cell type (**B**). Error bars indicate ± SEM, **** P < 0.0001, n = 5. Statistical significance was determined with an unpaired two-tailed T-test. Data are representative of two independent experiments.

**Fig. 4.**
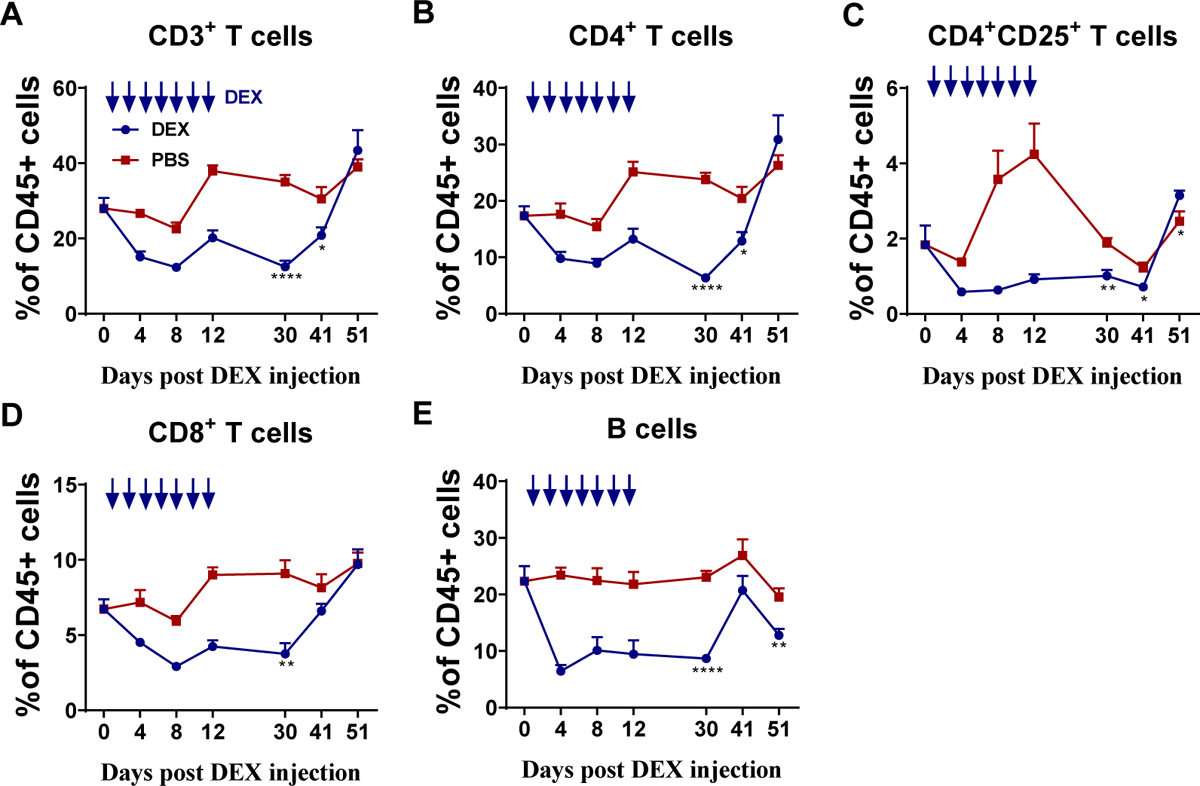
Recovery curves of lymphatic cells after drug withdrawal PBMC were isolated on days 9, 20, and 30 after DEX withdrawal and T cell subsets in CD45^+^ cells analyzed by flow cytometry. Antibody to B220 was used to identify B cells. Results from the DEX treatment group are shown in blue and from PBS vehicle group in red. Shown are the percentages of CD3^+^T cells (**A)**, CD4^+^T cells (**B)**, CD4^+^CD25^+^T cells(**C)**, CD8^+^T cells(**D)**, B220^+^B cells (**E)**. The results are presented as percentages in CD45^+^ cells. Error bars indicate ± SEM, **** P < 0.0001. Statistical significance was determined with an unpaired two-tailed T-test. Data are representative of two independent experiments.

### Immunosuppressive Effects Are Related to Dose of Dexamethasone

DEX is used in a wide range of doses to reduce the side effects of chemotherapy in patients with advanced cancer[7]. To gain insight into the immunoregulatory effects of DEX on the immune system, we selected high (5 mg/kg) and low (1.25 mg/kg) doses for study in Balb/c mice. We observed that quantities of the immune cells were reduced less by the lower dose of DEX (Figs. 5A, 5D, and sFig. 3A). However, macrophages and NKT cells were increased under the low dose of DEX (Figs. 5B-5D, sFigs. 3A, and 3B).

**Fig. 5.**
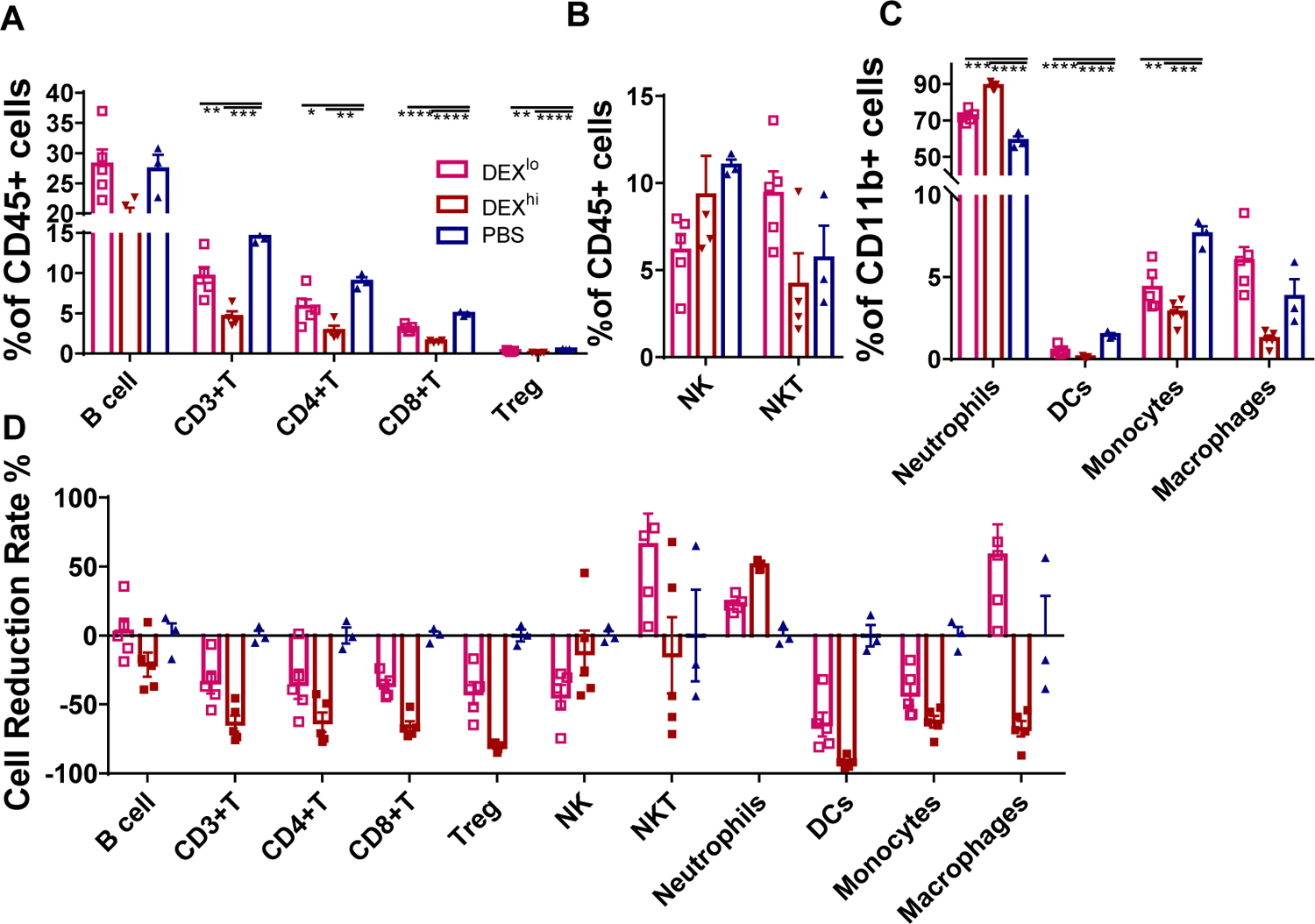
Immunosuppressive effects under different doses of DEX PBMC were isolated on day 8 after treatment with different doses of DEX and analyzed by flow cytometry. Antibodies to Ly6G, CD11c, F4/80, Ly6C, CD49b, CD3, Foxp3, and B220 were used to identify neutrophils, DC cells, macrophages, monocytes, NK, T, Treg, and B cells, respectively. Antibodies to CD3 and CD49b were used to identify NKT cells. The results are presented as percentages in CD11b^+^ cells (**A**) and CD45^+^ cells (**B, C)** and the proportion of the decline in each immune cell type (D). Error bars indicate ± SEM, **** P < 0.0001. Statistical significance was determined with an unpaired one-way ANOVA test.

### Dexamethasone Mediates Immunosuppression by Binding GR With a High Affinity

Since glucocorticoid-mediated immune modulation is attributed to glucocorticoid receptor-induced alterations in gene expressions, we investigated whether the immunoregulatory effects of DEX were related to GR signaling. Mifepristone (RU486), a progesterone receptor (PR) and GR antagonist that binds to GR with a low affinity, was given intraperitoneally every 12 h before each DEX administration. PBMC were collected 72 h after those treatments (sFigs. 3A-3B and 4A). We observed that RU486, as a GR-blocker, abolished the immune regulation by low-dose DEX. The proportion of T and NK cells returned to normal level (Fig. 6A). When a higher dose of DEX at 5 mg/kg was given, a more robust immunosuppressive function was obtained and there was less effect of RU486 (Fig. 6B). These data indicated that DEX was competitively bound to GR with RU486 and a stronger affinity was evident when given at higher dose.

**Fig. 6.**
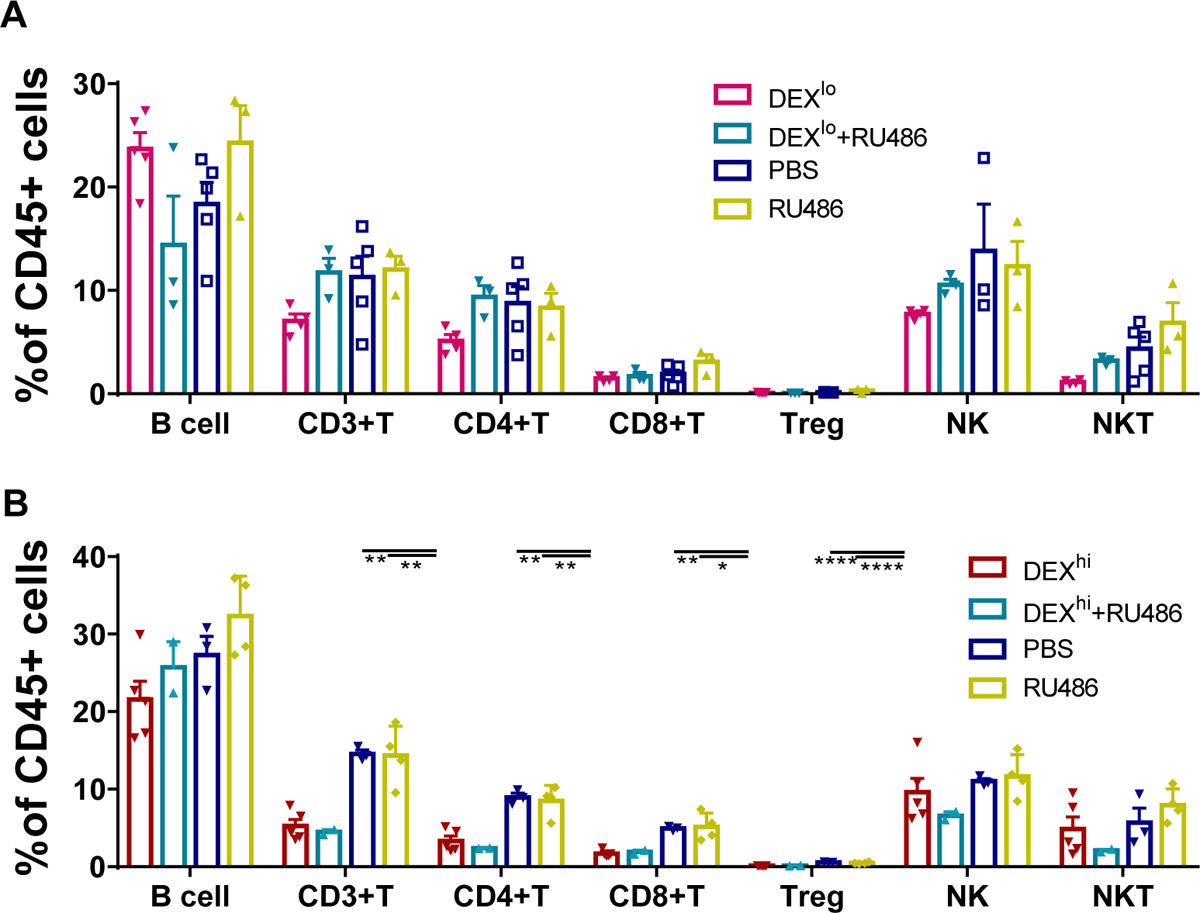
The high-affinity binding of DEX to GR reduces immune cells PBMC were isolated and analyzed by flow cytometry 3 days after treatment with 1.25 mg/kg DEX (**A**) or 5 mg/kg DEX (**B**). Antibodies to B220, Foxp3, and CD49b were used to identify B, Treg, and NK cells. Antibodies to CD3 and CD49b were used to identify NKT cells. The results are presented as percentages in CD45^+^ cells. Error bars indicate ± SEM, **** P < 0.0001. Statistical significance was determined with an unpaired one-way ANOVA test.

## Discussion

Our results demonstrated DEX’s immunoregulatory effects in down-regulating a wide range of immune cells in PBMC, both lymphoid and myeloid cell subsets. The doses of DEX in our study correspond to the doses used in clinical practice [19], suggesting that these findings may be relevant for understanding the immunosuppressive mechanisms of DEX in patients. Additionally, our data showed that the immunosuppressive function of DEX was dose-dependent, with higher doses leading to a stronger effect. This highlights the importance of carefully titrating the dosage of DEX in clinical settings to achieve the desired immunoregulatory outcomes. DEX, a synthetic glucocorticoid, had dramatic suppressive effects on DCs, monocytes, macrophages, NK, T, and B cells, but not on neutrophils and NKs.

Glucocorticoids signal through the glucocorticoid receptor (GR), a member of the superfamily of nuclear receptors that modulates both genomic and non-genomic regulatory pathways in almost every tissue in the bodies of mammals. Observed reductions of immune cells by the DEX treatments could be due to the fact that genomic regulation is receptor-dependent to suppress relevant gene expressions, and maybe also due to a mechanism of non-genomic regulation by inducing apoptosis [20, 21]. It has been reported that glucocorticoids could induce differential biological effects in dendritic cells based on GR isoforms to attenuate DC activity and maturation in one way [22] and in another to promote the differentiation of ‘tolerogenic’ DCs. Tolerance follows from down-regulation of the expression of MHC class II, co-stimulatory molecules (for example, CD80 and CD86) and pro-inflammatory cytokines (such as IL-12 and TNF) resulting in Treg induction [23]. Glucocorticoids also dampen T cell activation by interfering with TCR signaling and inhibition of interleukin (IL)-2 synthesis via binding to GR inside T cell cytoplasm has been reported[24]. Neutrophils are the earliest responding cell of the innate immune response to infections, particularly bacterial infections, and mount the initial inflammatory response.

They are also involved in the initiation of the inflammatory response by releasing pro-inflammatory cytokines and chemokines. Interestingly, we found that the proportion of neutrophils was increased by the DEX treatment, in direct contrast to the other immune cells. Observed enhanced neutrophils by the DEX treatments may achieve this by promoting the differentiation and activation of neutrophil progenitor cells in the bone marrow, leading to an increased production and release of mature neutrophils into circulation. The enhanced effect on the neutrophils could also be explained by recent studies in which neutrophils were found to be the cells more weakly responding to GCs and more resistant to DEX-induced apoptosis by mechanisms that included upregulation of anti-apoptotic proteins such as Mcl-1 and XIAP[25]. Alternatively, several studies have shown that neutrophils can also have immunosuppressive functions. They can suppress T cell proliferation and cytokine production, as well as promote the differentiation of Tregs[26, 27]. This dual role of neutrophils highlights their importance in maintaining immune homeostasis and preventing excessive inflammation.

Regarding B cells, immature B cells are more sensitive to glucocorticoid-induced apoptosis than mature B cells[15, 28], suggesting a regulatory role for glucocorticoids in B cell development. Additionally, glucocorticoids can also modulate B cell activation, proliferation, and antibody production. This highlights the complex interplay between neutrophils, B cells, and glucocorticoids in immune regulation and the delicate balance required for maintaining a properly functioning immune system, but more details of the effects of GCs on humoral immunity are needed.

Although there have been many studies and clinical reports of glucocorticoid effects on immunity during the preceding 60 years, this study has focused on the landscape of immunosuppression as contributed by key immune cell types, particularly those with roles in adaptive immunity. To our surprise, we observed that low DEX doses could increase the ratio of macrophages, while a reduction of macrophages was achieved when DEX was at a high dose. This double-edged sword effect of GCs on macrophages had not been reported previously and suggests a complex regulation via the GR signaling: at a low concentration, GC was reported to positively regulate NLRP3, a component of the inflammasome complex in macrophages, and thereby augment the pro-inflammatory response[29]. An alternative explanation might be that low GC concentrations, in the nanomolar range, exert an immunostimulatory effect on a range of macrophage functions, including adhesion, chemotaxis, phagocytosis, and cytokine production[30]. The macrophage is a crucial bridging arm from innate to adaptive immune activation and expresses pattern-recognition receptors, including Toll-like receptors that sense infectious agents and harmful signals, hence activating the inflammasome complex and secretion of inflammatory cytokines[31]. This cascade of reactivities serves to elicit adaptive immune responses.

Our findings confirm that DEX could depress immune cells via a high-affinity binding of GR but also show that this could be reversed to a normal level when using RU486 to block the GR. Treating severe cases of COVID-19 by duration with low doses of Dexamethasone (DEX) has displayed promising outcomes in terms of reducing inflammation and enhancing patient outcomes. Nonetheless, it is crucial to exercise caution when employing DEX due to its potential to exert immunosuppressive effects, which could potentially impede the body’s capability to mount an efficient immune response against the virus and secondary infections. Consequently, further investigation is warranted to ascertain the most optimal dosage and duration of DEX treatment in COVID-19 patients, with the objective of striking a delicate balance between curbing excessive inflammation and preserving an active immune response.

In conclusion, our study shows that treatment with a high dose of DEX negatively affects the numbers of both myeloid and lymphoid cells, except for neutrophils and NKT. Although myeloid cells returned to normal within 5 days of withdrawing the DEX treatment, the treatment affected lymphoid cells profoundly, and they took longer to recover. These findings will facilitate further understanding of the complexity of GCs effects on the immune responses and lead to a more economical use of the drug to achieve broad immune suppression.

## Author contributions

Bin Wang and Shuting Wu conceived, directed the experiments with input from all authors. Material preparation, data collection and analysis were performed by Shuting Wu. Shuting Wu, Shushu Zhao, Gan Zhao and Yiwei Zhong performed data analysis and interpretation. The first draft of the manuscript was written by Shuting Wu and all authors commented on previous versions of the manuscript. All authors read and approved the final manuscript.

## Data Availability

The data generated in this study are available upon request from the corresponding author.

## Ethics approval

This study was performed in line with the guidelines of the Institutional Animal Care and Use Committee of the Fudan University in China. All experiments were approved by the Committee of Experimental Animals of SHMC with the reference number 202012037S.

## Abbreviations

DCs: dendritic cells

DEX: dexamethasone

DMSO: dimethyl sulfoxide

FCM: flow cytometry

GC: Glucocorticoid

GR: glucocorticoid receptor

HPA: hypothalamic-pituitary–adrenal

*ip*: intraperitoneal

MHC: major histocompatibility complex

PBMC: peripheral blood mononuclear cells

PBS: phosphate-buffered saline

PR: progesterone receptor

RU486: Mifepristone

TCR: T-cell receptor

