## Supplemental Figures for "Revisit the Inhibitory Effects of Glucocorticoids on Immunocytes"

### SUPPLEMENTARY FIGURES

#### sFigure 1.

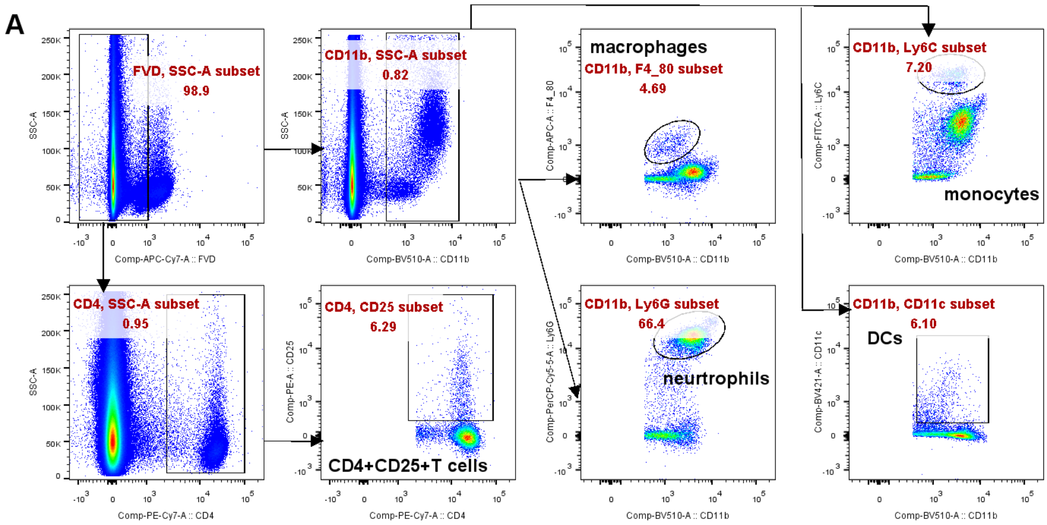

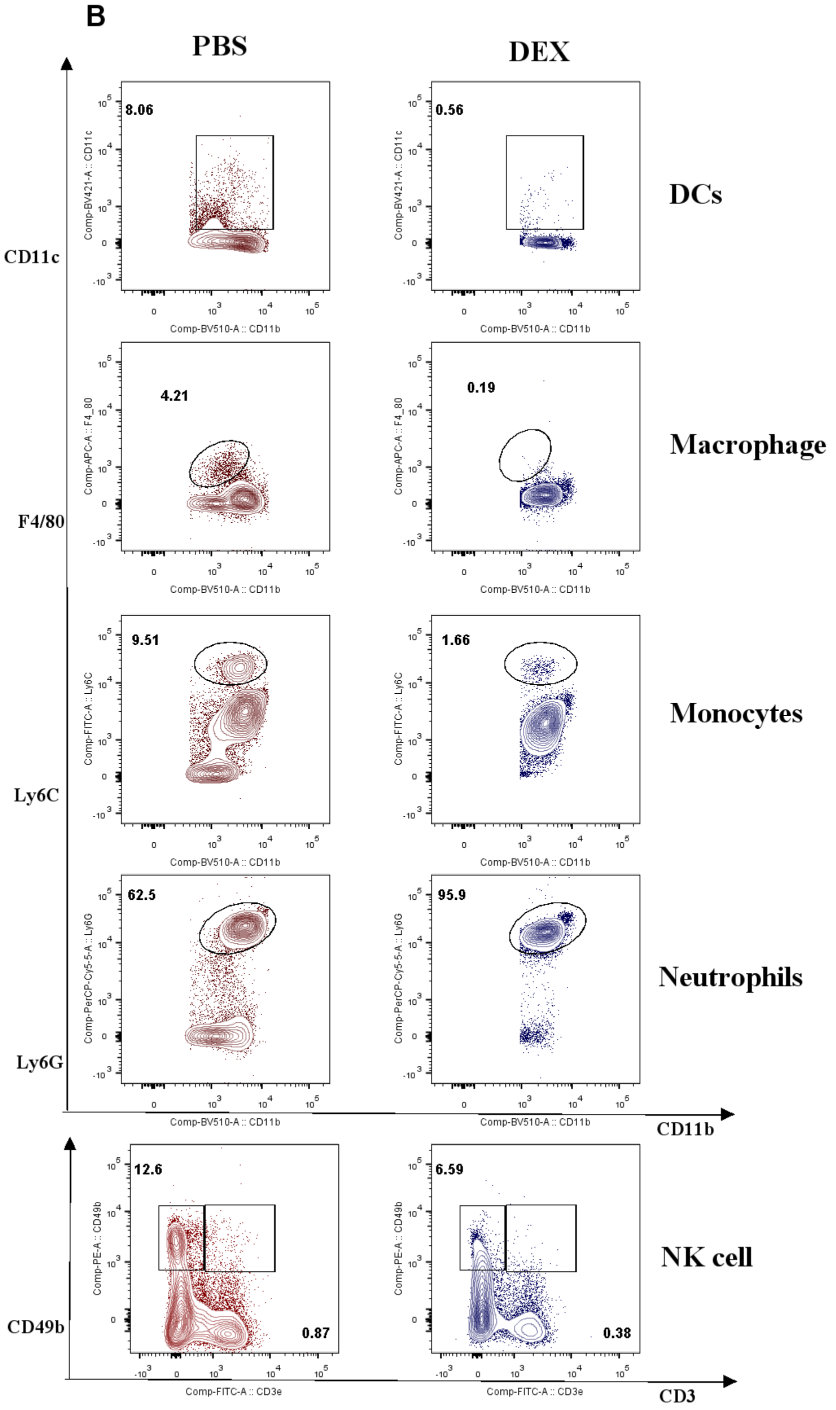


**sFigure. 1** Gating Strategies for identification of myeloid-derived cells subsets

(A). Experimental outline and gating strategy for identification of myeloid-derived innate immune cells subsets within PBMC of treatment mice by flow cytometry. (B). Representative data between DEX treated mice and PBS treated mice were described by flow cytometry.

sFigure 2.

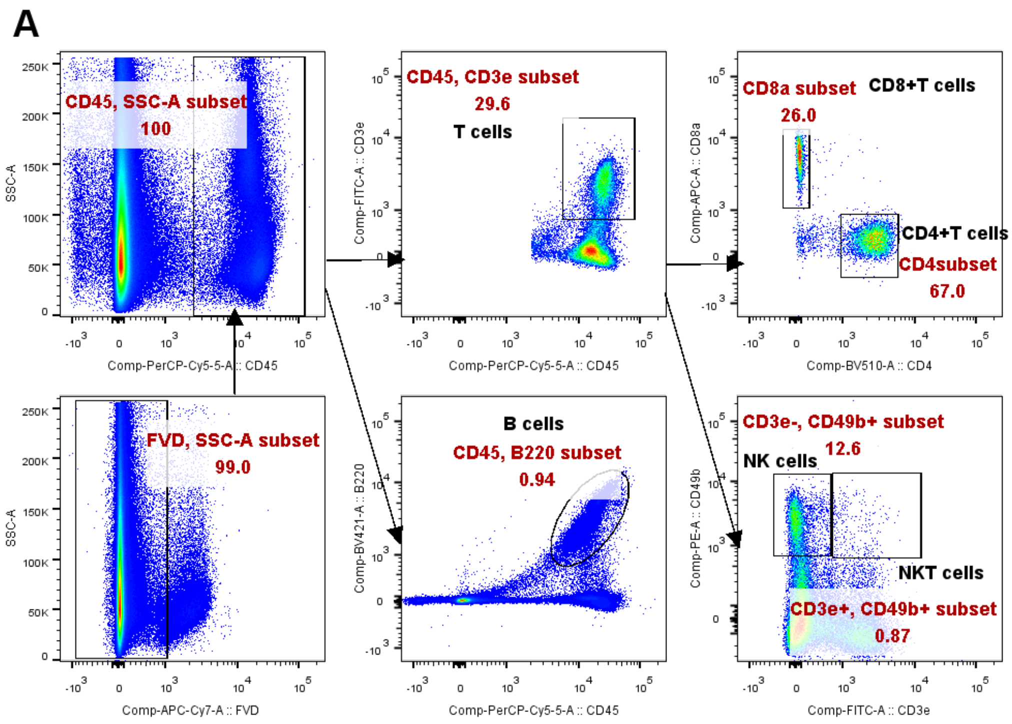
A
CD45, SSC-A subset
100
FVD, SSC-A subset
99.0
CD45, CD3e subset
29.6
T cells
B cells
CD45, B220 subset
0.94
CD8a subset
26.0
CD8+T cells
CD4+T cells
CD4subset
67.0
CD3e-, CD49b+ subset
12.6
NK cells
NKT cells
CD3e+, CD49b+ subset
0.87
SSC-A
Comp-PerCP-Cy5-5-A :: CD45
Comp-APC-Cy7-A :: FVD
Comp-FITC-A :: CD3e
Comp-BV421-A :: B220
Comp-APC-A :: CD8a
Comp-BV510-A :: CD4
Comp-PE-A :: CD49b

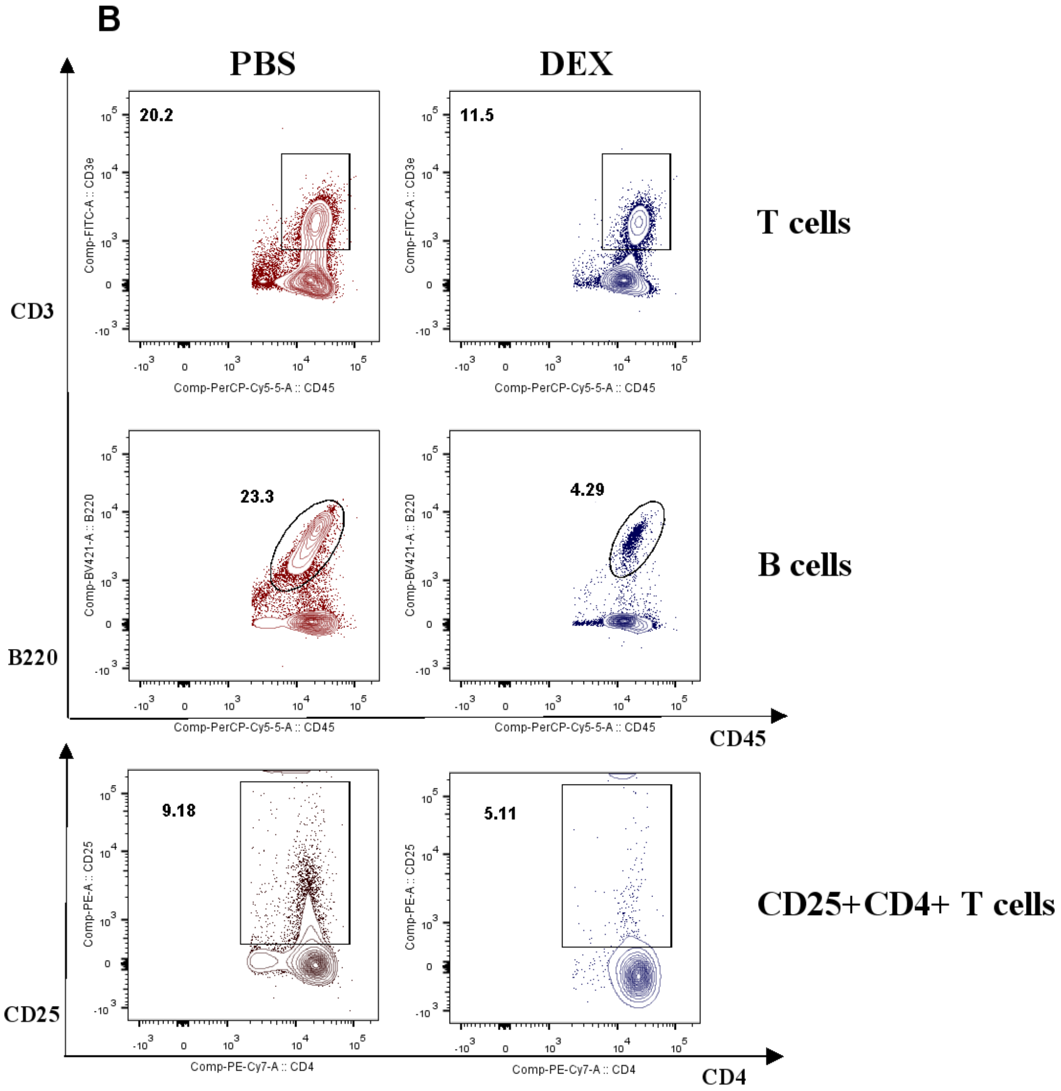


**sFigure. 2** Gating Strategies for identification of lymphoid-derived cells subsets

(A). Experimental outline and gating strategy for identification of lymphoid-derived cells subsets within PBMC of treatment mice by flow cytometry. (B). Representative data between DEX treated mice and PBS treated mice were described by flow cytometry.

**sFigure 3.**

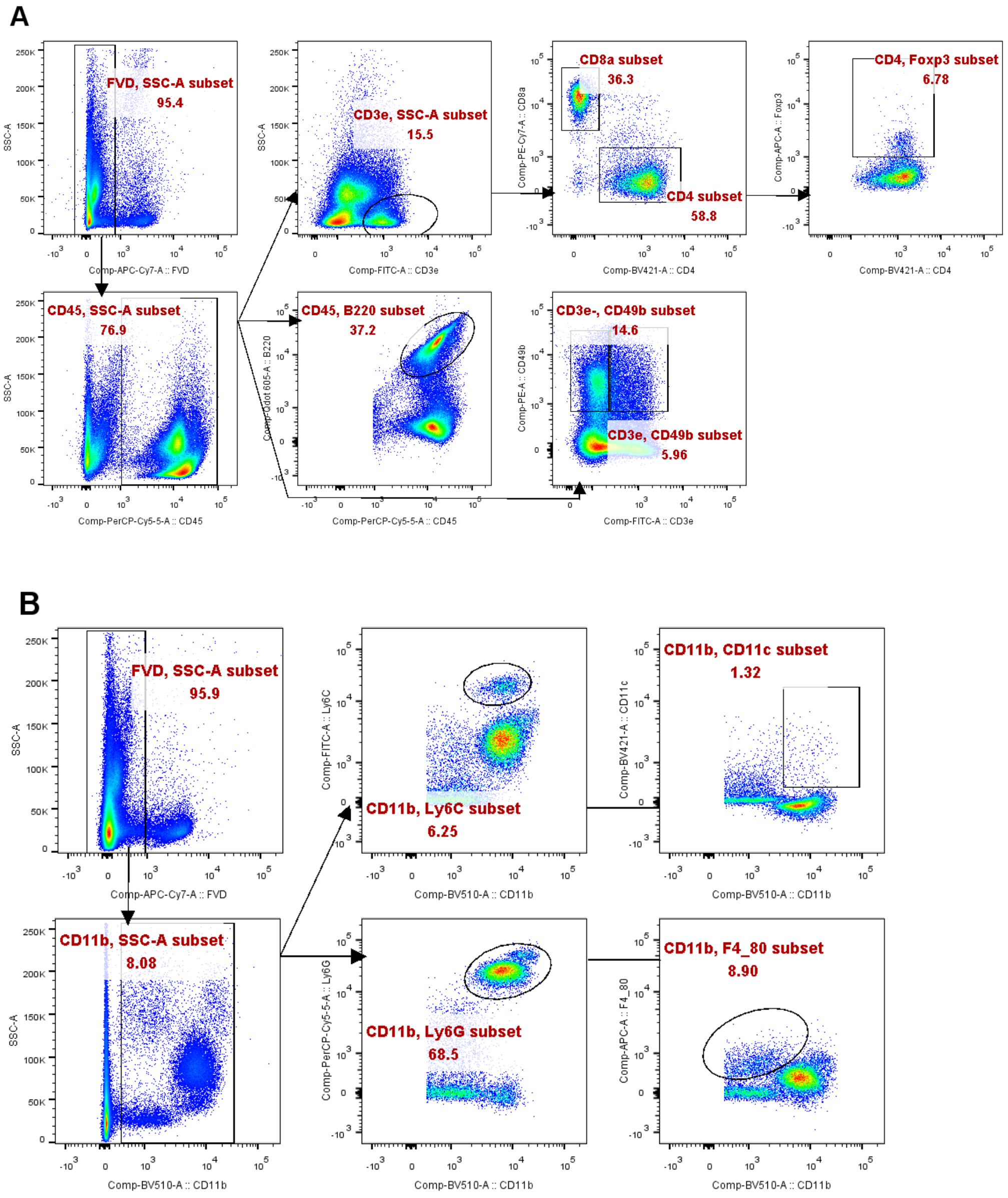


**sFigure. 3** Gating Strategies for identification of myeloid-derived and lymphoid-derived cells subsets

(A). Experimental outline and gating strategy for identification of lymphoid-derived cells subsets within PBMC of treatment mice by flow cytometry. (B). Experimental outline and gating strategy for identification of myeloid-derived cells subsets within PBMC of treatment mice by flow cytometry.

**sFigure 4.**

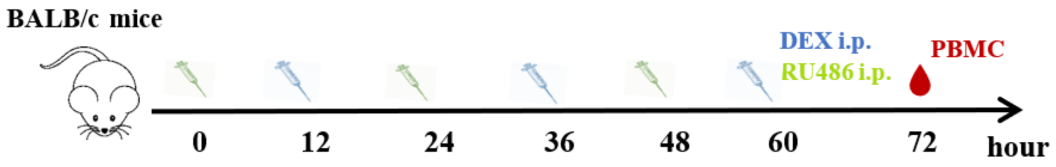


**sFigure. 4** Administration diagram

Schematic diagram of DEX and RU486 competitively binding GR.
